# Assessment of genetic structure of the endangered forest species *Boswellia serrata* Roxb. population in central india

**DOI:** 10.1101/412874

**Authors:** Vivek Vaishnav, Shashank Mahesh, Pramod Kumar

## Abstract

*Boswellia serrata* Roxb., a commercially important species for its pulp and pharmaceutical properties was sampled from three locations representing its natural distribution in central India for genetic characterization through 56 RAPD + 42 ISSR loci. The wood fiber dimensions measured for morphometric characterization confirmed 11.36% of the variation in the length and 8.75% of the variation in the width indicating its fitness for local adaptation. Bayesian and non-Bayesian approach based diversity measures resulted moderate within population gene diversity (0.26±0.17), Shannon’s information index (0.40±0.22) and panmictic heterozygosity (0.28±0.01). A high estimate for genetic differentiation measures i.e. G_ST_ (0.31), G_ST_-B (0.33±0.02) and θ-II (0.45) led to the distinct clusters of the sampled genotypes representing their regional variability due to limited gene flow and total absence of natural regeneration. We report the first investigation of the species for its molecular characterization emphasizing the urgent need for the genetic improvement program for the *In-situ*/*Ex-situ* conservation and sustainable commercialization.

## INTRODUCTION

*Boswellia serrata* Roxb. (family-Burseraceae), commonly known as salai guggul or Indian frankincense (olibanum indicum) is a commercially important deciduous tree of India (Shah et al. 2008). It is found in the region having rainfall between 500mm-2000mm and temperature up to 45 °C. The species has the ability to thrive in the poorest and the shallowest soils (Bhat et al. 1952). In India, it is distributed in states Rajasthan, Maharashtra, Madhya Pradesh, Karnataka and Chhattisgarh (Pawar et al. 2012). The oleo-gum-resin of the tree contains boswellic acid, which is effective for the treatment of inflammatory disorders, arthritis, cardiovascular diseases (Ammon 2006), diarrhea, dysentery and other skin diseases (Khare 2004). Apart from these medicinal values, the resin of the species was found a more effective sizing agent in papermaking than the rosin obtained from *Pinus* species (Sharma et al. 1985). The convincing wood quality for pulp making led to the establishment of the first paper mill in India in Nepanagar of Madhya Pradesh (Khan 1972) as the region had been occupied by wide range of natural patch of the species.

The species is highly out-crossing supported by the self-incompatibility to selfing (Sunnichan et al. 2005). But, a poor fruit setting (2.6% - 10%) under open pollination condition, inadequate production of viable seeds and scanty seed germination (10% - 20%) limit the distribution of the species in nature (Ghorpade et al. 2010). The species population has been harvested for the frankincence also (Sharma 1983). The scarcity of protocols to regenerate it through seeds and clones makes the mass multiplication difficult (Purohit et al. 1995). This situation resulted in declined abundance of the species and the International Union for Conservation of Nature (IUCN) has enlisted it under the status of endangered. Therefore, the available natural patches of the species require keen attention for both *In-situ* and *Ex-situ* conservation with actual knowledge of available genetic resources of the species for management and breeding objectives.

Information on actual genetic variability and a cryptic number of the genetically differentiated genetic resource of any species is an important aid for its conservation and genetic improvement. For such purposes, DNA based markers are of value because unlike the morphometric traits the molecular markers are independent of the influence of environment and can be assessed at any growth stage. Among the various molecular markers employed to assess diversity studies, PCR-based dominant markers such as RAPD (random amplified polymorphic DNA, Williams et al. 1990) and ISSR (inter-simple sequence repeat, Zietkiewiez et al. 1994) have become popular due to its polymorphism and discrimination power, as their application does not need any prior sequence information. These primer systems have been successfully applied for the genetic characterization of populations of tropical tree species (Ansari et al. 2012, Abuduli et al. 2016, Vaishnav & Ansari 2018). Bayesian statistics has been extended to dominant markers for precise estimates of population genetic hierarchies equivalent to those obtained with codominant markers, circumventing inbreeding estimate within the population (Holsinger et al. 2002). Therefore, the present investigation was conducted to differentiate the morphometric and the genetic variability exhibited by the natural population of *B. serrata* Roxb. in three agro-climatic regions of central India applying dominant marker system.

## MATERIAL AND METHODS

### Population sampling

The information of the distribution of the patches or population of the species was obtained from the old forest working plans of the concerned forest departments of the Indian state Chhattisgarh, which is classified in three agro-climatic zones *viz.* Northern Hills, Chhattisgarh plain and Bastar plateau (Table 1).

**Table 1.** Geo-climatic conditions of the three regions of natural distribution of *B. serrata* Roxb. populations sampled for the investigation

| Regions | Agro-climatic zones | GPS coordinate ranges | | Altitude (m) | Mean annual temperature ($^{\circ}$C) | | Annual Precipitation (mm) |
| --- | --- | --- | --- | --- | --- | --- | --- |
|  |  | Latitude (N) | Longitude (E) |  | Min | Max |  |
| Dhamtari (DT) | Chhattisgarh Plain | 20.37861-20.40511 | 81.96636-81.98122 | 445.80 ±5.44 | 21.60 ±5.22 | 33.02 ±3.56 | 1488±0 |
| Narayanpur (NP) | Bastar Plateau | 19.71365-19.71586 | 81.19447-81.19836 | 595.60 ±21.39 | 20.80 ±5.22 | 32.40 ±3.36 | 1628±0 |
| Sarguja (SG) | Northern Hills | 23.39300-23.39446 | 83.46277-83.46783 | 619.75 ±7.15 | 19.02 ±6.35 | 31.02 ±4.07 | 1670±0 |

± - Standard deviation (SD)

Forests of these agro-climatic regions were visited during July 2014 to September 2014 to collect samples from 20 trees of each region in natural distribution of *B. serrata* respecting inter-tree distance of 100 m along the latitude (Figure 1).

**Figure 1.**
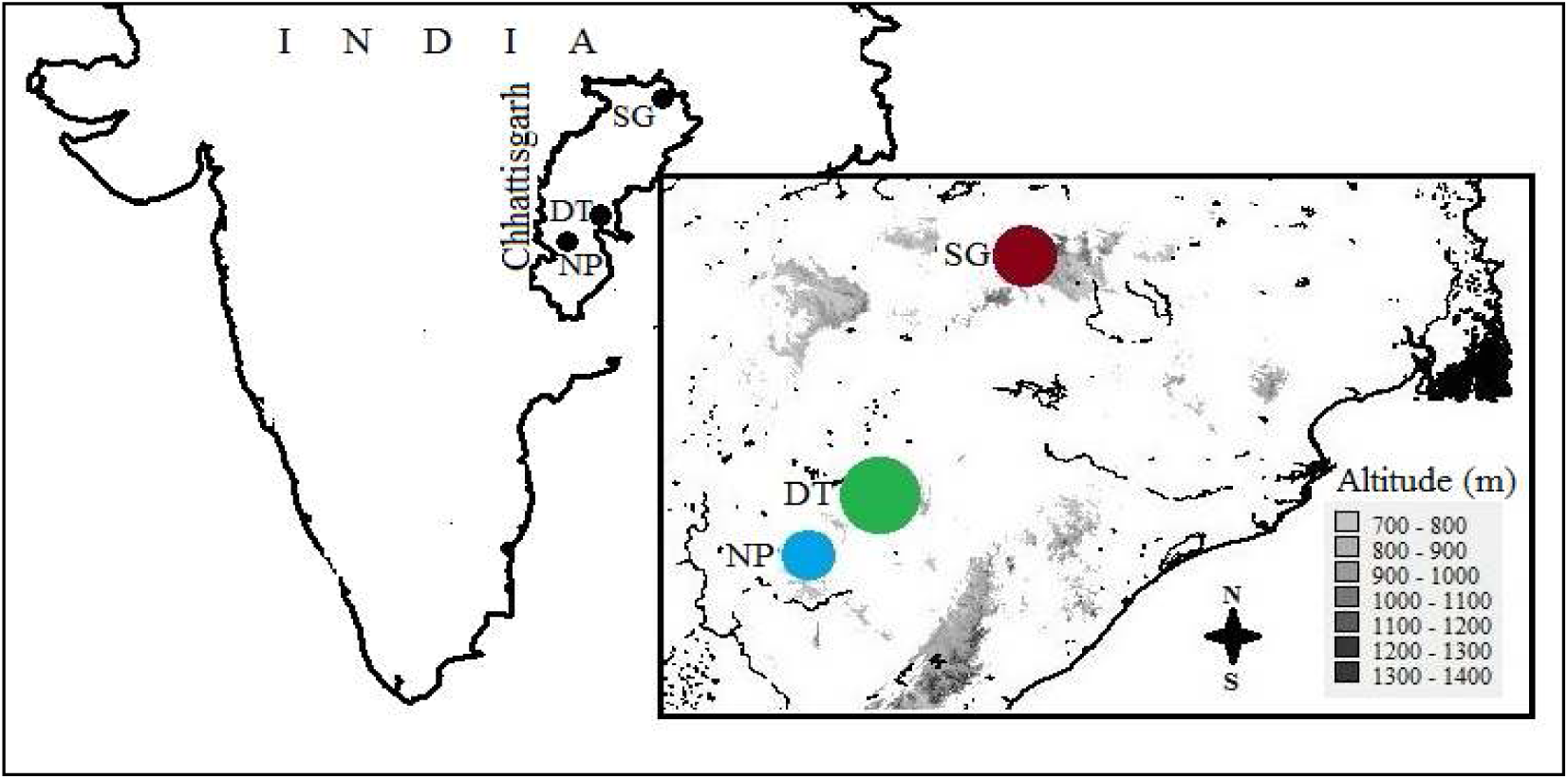
Three regions i.e. Sarguja (SG), Dhamatri (DT) and Narayanpur (NP) of natural distribution of *B. serrata* Roxb from central Indian region (Chhattisgarh) were sampled for the investigation. Every location represents distinct agro-climatic zone namely Northern hills (SG), Chhattisgarh plain (DT) and Bastar plateau (NP). An inset view of altitudinal variation is given in the box and size differences among the location’s legend dot depicts differences in gene diversity of the species estimated employing the dominant markers

The girth at breast height (GBH) was measured by measuring tape. A wood radial core sample was extracted from each tree at breast height (1.34 m) and was stored in a tube with 40% formaldehyde. A leaf sample was also collected from each tree. The samples were brought to the laboratory in a cryo-box.

### Measurement of wood fiber dimensions

In the laboratory, the wood samples were macerated following the method described in Mahesh et al. (2015). Five slides were prepared for each tree and the wood fiber length (WFL) and width (WFW) of five fibers per slide were measured through the compatible program integrated with the compound microscope (Leica, EC3, Switzerland).

### DNA isolation and amplification

DNA from the leaf samples was isolated following modified CTAB method (Deshmukh et al. 2007). To select the reproducible markers, 10 RAPD and 10 ISSR primers were cross-amplified on set of 12 genotypes of the species (four from each region) and finally, only five from each of RAPD and ISSR markers were selected based on their polymorphism and reproducibility for the final amplification of all sixty genotypes (Table 3). For RAPD/ISSR amplification, the final reaction mixture for PCR assay was consisted of 15 µl/10 µl containing 50 ng/ 40ng genomic DNA, 0.66 µM/0.8 µM of primers, 0.2 mM/ 0.1mM of dNTP mix, 1.5 mM/2.5 mM MgCl_2_, 1X buffer with KCl and 1 unit of Taq polymerase. PCR parameters included an initial 4 minutes denaturation step at 94 °C followed by 35 cycles of 30s at 94 °C, 30s at the annealing temperature of 35 °C/50 °C, 45s extension at 72 °C followed by a final extension of 5 minutes at 72 °C. The amplified products were electrophoresed on 1.5% agarose gel containing 0.5µg/ml ethidium bromide (EtBr) in 0.5 X TBE (pH 8.0). The separation was carried out by applying constant 100V for 3 hours. The fractionated amplified products on agarose gel were visualized on gel documentation system under UV light. To avoid the homology, the amplification profile of all sixty genotypes was evaluated through the molecular weight of the bands for each primer and the bands were scored in a Microsoft Excel sheet in binary data format.

### Data analysis

The measures of central tendency and the coefficient of variation (CV) were calculated for GBH, WFL, and WFW. A genetic profile of 60 genotypes on 10 markers was generated and analyzed following both band-based and allele frequency-based approaches as suggested in Bonin et al. (2007). The information from the markers was evaluated estimating mean allelic frequency (AF), gene diversity (GD), and polymorphic information content (PIC) by program POWERMARKER v3.1 (Liu & Muse 2005). Resolving power (RP) of the markers was calculated following the formula given by Prevost and Wilkinson (1999).

The genetic variability and the differentiation of the sampled population were evaluated applying both non-Bayesian and Bayesian approaches. The program POPGENE v1.32 (Yeh et al. 1999) was applied for the non-Bayesian estimates of genetic diversity measures i.e. Nei’s gene diversity (GD), Shannon’s information index (I) and the percentage of polymorphism (P%) for each location and G_ST_ was calculated to evaluate the level of genetic differentiation. The genetic relationship among the genotypes was calculated based on Jaccard’s genetic similarity coefficient by the program DARWIN v5.0 (Perrier & Jacquemoud-Collet 2006). The principal coordinate analysis (PCoA) was performed to cluster the genotypes in different axes applying program GENALEX v6.5 (Peakall & Smouse 2012). Analysis of molecular variance (AMOVA) was performed applying program ARLEQUIN v3.11 (Excoffier & Lischer 2010) to estimate the hierarchical variation. The isolation by distance (IBD) was confirmed calculating the correlation coefficient between two pairwise matrices of genetic and geographical distances among genotypes through Mantel’s test performed by program Alleles in Space (Miller 2005) in independent runs (with and without logarithmic transformation) on 10,000 permutations.

For the Bayesian approach based genetic estimates we preferred the θ-statistics (*viz.* θ-I, θ-II, θ-III and G_ST_-B) implemented in program HICKORY v1.1 (Holsinger & Lewis 2007) that allows direct estimation of genetic differentiation measure (F_ST_) from dominant markers without assumption of prior knowledge of the extent of inbreeding and Hardy –Weinberg Equilibrium (HWE) within population even with small size and number. Region-wise panmictic heterozygosity (Hs) and total panmictic heterozygosity (Ht) were also estimated. All these estimations were performed with 50,000 steps of burn-in, 500,000 replicates, ‘thinning’ = 20 following the best-suited model for the data amongst the all four models (*viz.* full, f = 0, θ = 0 and f-free models) implemented in the program based on the minimum DIC (deviance information criterion) value as suggested by Spiegelhalter et al. (2002). To determine the most suitable number of cryptic populations (K) for the dataset, the program STRUCTURE v2.3.1 (Pritchard et al. 2000) was run applying model combinations of admixture/ no-admixture model with correlated/ independent allele frequencies among the samples on 100000 burn-in and 1000000 MCMC repeats with three-run from K=1 to K=9, for number of K 2≤ K ≤ 8. The most suitable model and the best-suited ‘K’ was determined based on the highest Delta-K value resulted by online program STRUCTURE HARVESTER (Earl and VonHoldt 2012).

## RESULTS

### Variation in wood fiber dimension

The WFL of the species ranged between 0.803 mm to 1.397 mm with average of 0.968±0.11 mm (11.36% CV) and the WFW of the species ranged between 0.019 mm to 0.030 mm with average of 0.025±0.002 mm (8.75% CV). Both traits followed normal distribution and no significant correlation was observed between them. WFL and WFW both were evenly distributed on the standard deviation (σ) based classes and no significant variation was observed among the locations based on WFL and WFW (Table 2).

**Table 2.** Variability in wood trait parameters of three sub-populations of *B. serrata* Roxb.

| Regions | GBH<br>(m) | WFL<br>(mm) | WFW<br>(mm) |
| --- | --- | --- | --- |
| Dhamtari<br>(DT) | 0.99<br>±0.25 | 0.919<br>±0.097 | 0.025<br>±0.001 |
| Narayanpur<br>(NP) | 1.07<br>±0.22 | 0.975<br>±0.095 | 0.026<br>±0.002 |
| Sarguja<br>(SG) | 1.40<br>±0.25 | 1.009<br>±0.120 | 0.024<br>±0.003 |
| Average | 1.15<br>±0.30 | 0.968<br>±0.11 | 0.025<br>±0.002 |
| CV (%) | 26.01 | 11.36 | 8.75 |

GBH- girth at breast height, WFL- wood fiber length, WFW- wood fiber width, CV- coefficient of variation, ±- standard deviation (SD)

### Genetic informativeness of markers

The five RAPD primers amplified 826 bands and 56 loci (14.75 bands/locus). The diversity measures AF and GD were 0.81±0.065 and 0.27±0.075 respectively. PIC and RP were 0.22±0.054 and 5.51±3.17 subsequently (Table 3).

**Table 3.**
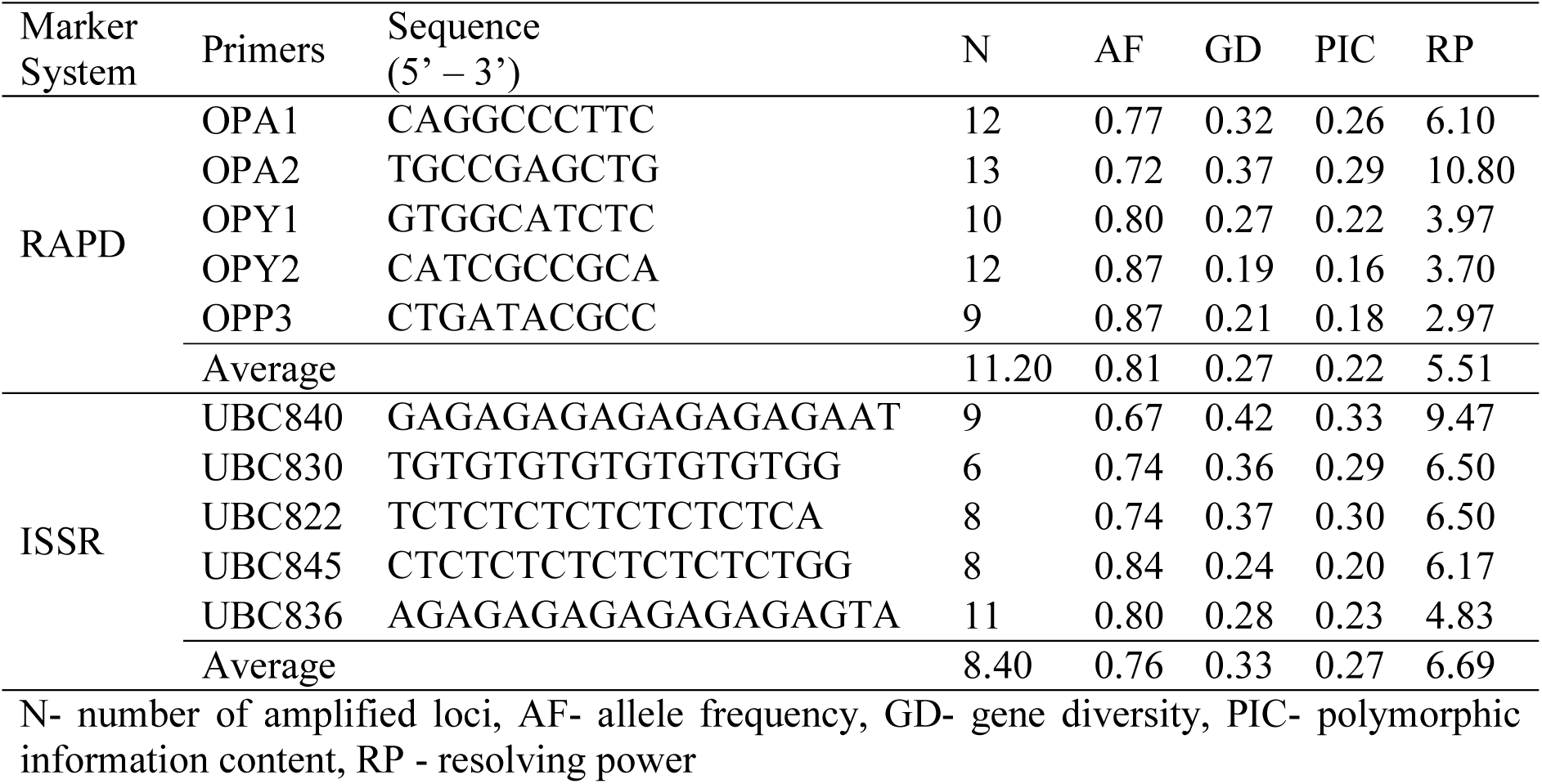
Genetic informativeness of RAPD and ISSR markers

For RAPD markers, the diversity measures and discriminatory power measures were found in significant (p<0.05) linear correlation. The five ISSR primers amplified 1004 bands for 42 loci (23.90 bands/locus). In a significant linear correlation (p<0.01) to each other, the AF was 0.76±0.068 and the GD was 0.33 ±0.073. The PIC and the RP were 0.27 ±0.052 and 6.69±1.69 subsequently (Table 3).

### Genetic diversity and differentiation

Among the non-Bayesian estimates of genetic diversity measure for the species population, 100% polymorphism, 0.26±0.17 GD, and 0.40±0.22 I were observed among the regional locations (Table 4). For all measures, no significant difference was found among the three regions. Dhamtari was found slightly more diverse and polymorphic than the other two regions (Table 4). The G_ST_ was 0.31. The Bayesian model implemented in Hickory estimated the lowest DIC value (896.96) for the full model and the panmictic heterozygosity was 0.28±0.01. No significant variation in panmictic heterozygosity was found among the three regional sub-populations (Table 4). The θ-I, θ-II, and θ-III values were 0.50±0.04, 0.45±0.04 and 0.25±0.01 subsequently and the G_ST_-B was 0.33±0.02.

**Table 4.** Population genetics parameters for 60 genotypes sampled from three sub-populations of *B. serrata* Roxb.

| Regions | P% | GD | I | Hs |
| --- | --- | --- | --- | --- |
| Dhamtari (DT) | 63.27 | 0.20<br>±0.20 | 0.30<br>±0.28 | 0.20<br>±0.01 |
| Narayanpur (NP) | 57.14 | 0.19<br>±0.19 | 0.28<br>±0.28 | 0.19<br>±0.01 |
| Sarguja (SG) | 55.10 | 0.14<br>±0.16 | 0.22<br>±0.24 | 0.17<br>±0.01 |
| Overall* | 100 | 0.26<br>±0.17 | 0.40<br>±0.22 | 0.28<br>±0.01 |

± SD, P% - polymorphism%, GD - Nei (1973) gene diversity, I - Shannon's information index, Hs - panmictic heterozygosity resulted by Hickory, \* considering all 60 genotypes as a group

The dendrograph based on Jaccard’s genetic similarity coefficient (average 0.67) grouped the genotypes following their regions (Figure 2). The genotypes sampled from Sarguja further bifurcated into two different but conjoint clusters supported by the bootstrap values >60 (Figure 2).

**Figure 2.**
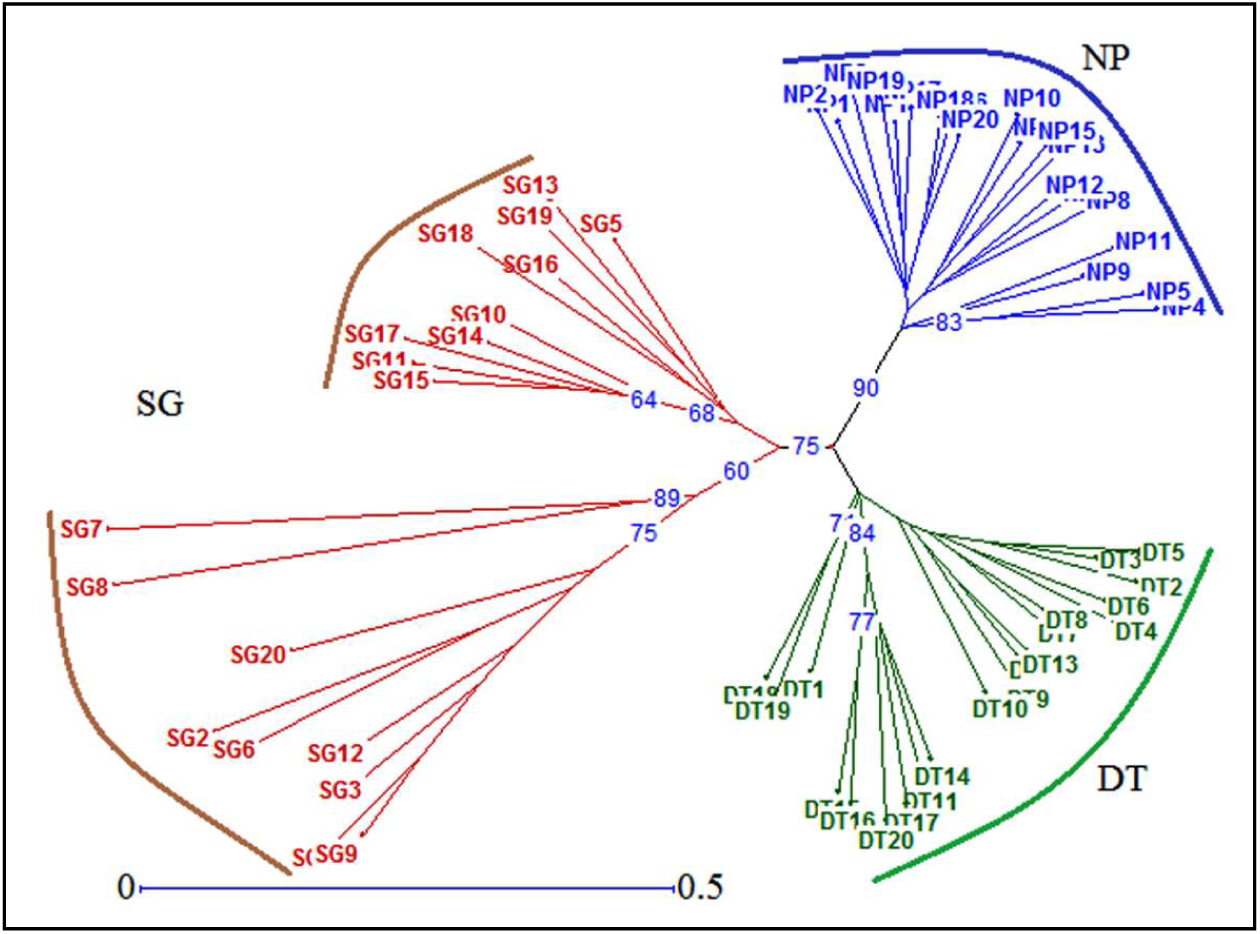
A Jaccard’s similarity coefficient based radial dendrograph of 60 accessions from three natural populations i.e. Sarguja (SG), Dhamtari (DT), and Narayanpur (NP) of *B. serrata* Roxb. Bootstrap values (>60) are shown on nodes of accessions

The PCoA accounting for 68.97% cumulative separation based on the same genetic distance separated the genotypes of Narayanpur from rest of the sampled genotypes and the equal proportion of the genotypes from Dhamatari and Sarguja was in the separate group from their sampled locations making overall four small clusters (Figure 3).

**Figure 3.**
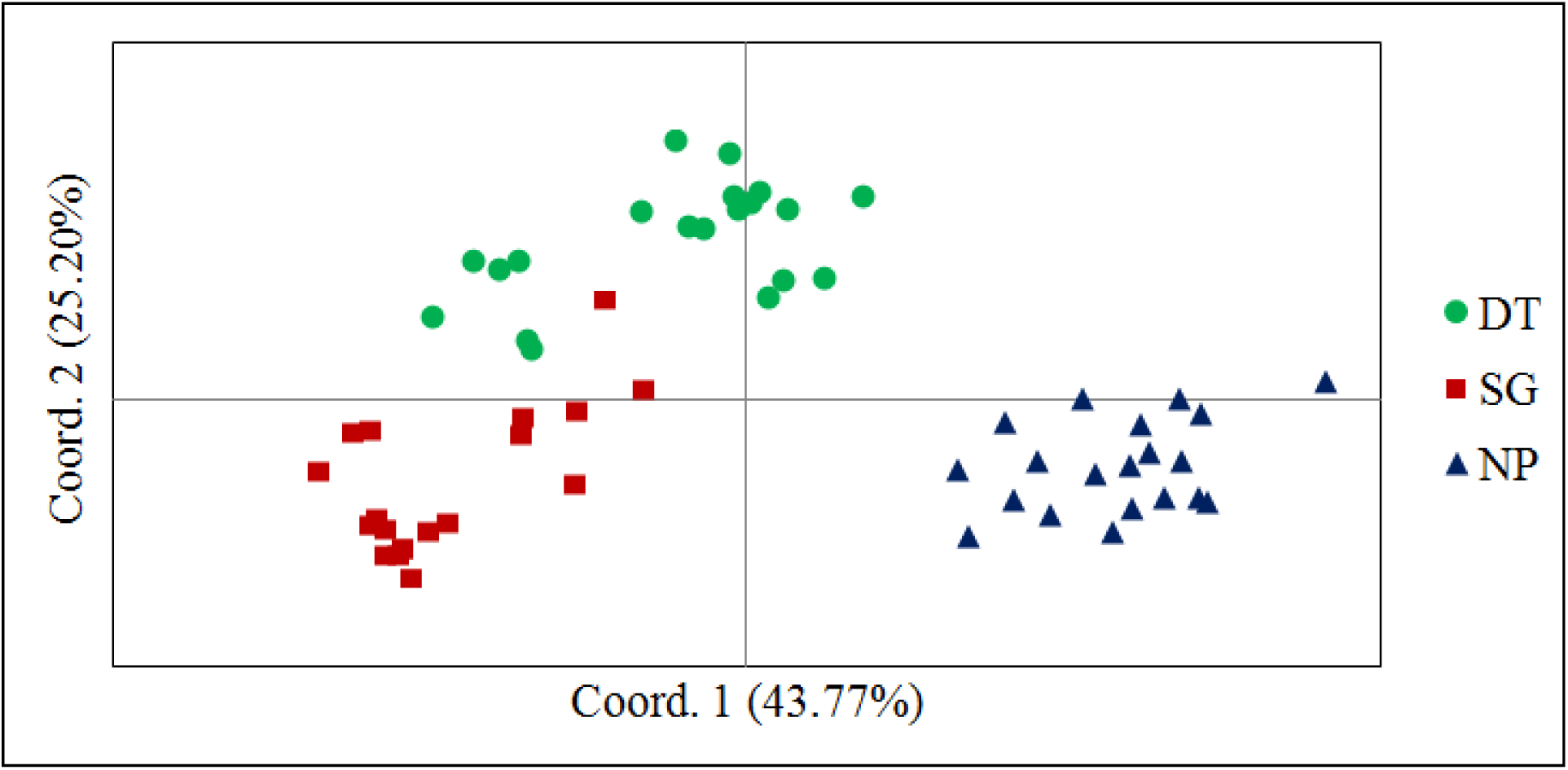
The principal coordinate analysis, showing closeness of genotypes of *B. serrata* Roxb sampled from Dhamtari (DT) and Sarguja (SG) and the genotypes from Narayanpur (NP) cluster distinctly, in two coordinates covering 68.97% of variation

The Bayesian program STRUCTURE assigned K=2 as the most appropriate number of cryptic population for the sampled genotypes based on the highest deltaK value in STRUCTURE HARVESTER. The bar plot generated (Figure 4) shows admixing of genotypes assigned to the two cryptic populations (K=2) with >80% of ancestry coefficient. The genotypes from Dhamtari were found admixed with both of the cryptic populations, the genotypes from Narayanpur and Dhamatri were assigned to be in the same cluster. On the other hand, genotypes from Sarguja clustered distinctly (Figure 4). AMOVA resulted 46.03±0.65% variation among groupings suggested by dendrograph, PCoA, and STRUTURE (K=2) and 53.38±0.15% variation within these groups. Mantel’s test found significant (p<0.001) correlation between pair-wise genetic and geographical distances among the genotypes.

**Figure 4.**
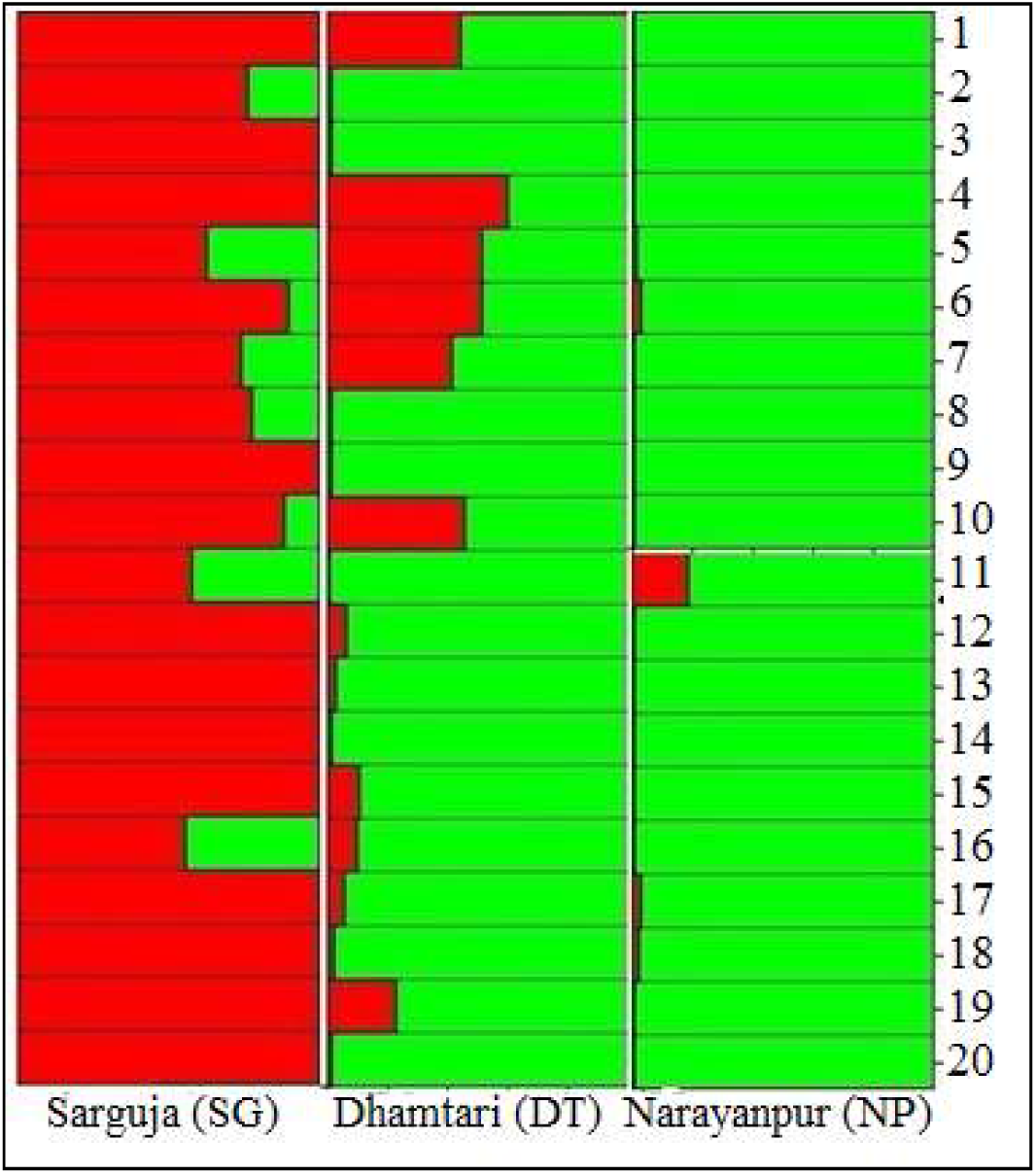
The Bayesian clustering resulted by program STRUCTURE depicted two cryptic populations (K) in 60 genotypes of *B. serrata* Roxb sampled from three locations of its natural distribution. Each genotype is presented by a separate colored bar with corresponding number. Green and Red color represents two different cryptic populations.

## DISCUSSION

*B. serrata* Roxb. a member of family Burseraceae, bifurcated from “Terebinthaceae” along with Anacardiaceae during the Miocene epoch due to lesser intrinsic ability to adopt the changing conditions across the climatic niches (Weeks et al. 2014). Therefore, the distribution of genus *Boswellia* limited within the mediterranian and tropical regions. *B. serrata* Roxb. is found only in central India and the species is renowned for its olio-gum-resin and pulp content. Due to uncontrolled harvesting, it has become scarce in nature and the literature regarding its distribution is also not adequate to locate it. During the investigation, the species was found in the temperature range of 19.02±6.35 °C to 33.02±4.07 °C and the annual rainfall of 1595.33±95.30 mm confirming its suitability for moist conditions along with the arid regions. For molecular characterization of the species, the RAPD and ISSR markers are preferred due to the absence of background genomic information of the species to design microsatellites and low-cost applicability. The problems with the dominant markers such as low reproducibility and uncertain homology of alleles have been handled during amplification and data profiling. Bayesian model-based θ-statistics has been applied based on its successful estimation of the genetic variability and the differentiation of population applying dominant markers system (Holsinger & Wallace 2004) in forest species as well (Ci et al. 2008,Vaishnaw et al. 2015).

### Variation in wood fiber dimension

The wood fiber length along with the thickness is considered to be the most important characteristics for the mechanical strength of wood (Migneault et al. 2008) and for the making of high-quality groundwood in paper industries (Zobel & Jett 1995). The comparison with the other tropical trees employed in pulp making process establishes that the wood fiber dimensions of *B. serrata* Roxb. obtained in our investigation is much higher than those of *Ailanthus altissima* (WFL= 0.747±0.035 mm and WFW= 0.023±0.0004 mm) and *Eucalyptus globulus* (WFL = 0.785±0.012 mm and WFW= 0.018±0.0005) reported in Baptista et al. (2014). The WFL is lower than of *Eucalyptus grandis* (1.01±0.14 mm) reported in Bhat et al. (1990). The coefficient of variation of wood fiber dimension in *B. serrata* is higher than *A. altissima* (WFL= 4.76% and WFW= 1.66%) and *E. globulus* (WFL= 1.5% and WFW= 2.72%) and lower than *E. grandis* (WFL= 14.45%) in above studies. In terms of adaptability, such high variation confirms the adaptive fitness of the population to the local ecological conditions (MacPherson et al. 2015).

### Genetic informativeness of the primers

With an approach to handle the limitation of the non-reproducibility, the dominant markers (RAPD/ISSR) have exhibited moderate values for AF and GD. It indicates that the approach avoiding null alleles have controlled the informativeness of the primers helping genetic differentiation (Lynch & Milligan 1994). The AF exhibited by the RAPD/ISSR markers are slightly higher to the information resulted in populations of *Tectona grandis* L. f. sampled from India on ISSR markers (0.73±0.02, Vaishnav & Ansari 2018). On the other hand, negligible differences in AF among the markers of RAPD and ISSR systems ensure the qualitative support for the diversity assessment (Lynch & Milligan 1994, Nybom & Bartish 2000). The discriminatory power measures exhibit a strong and linear relationship between the ability of a primer to distinguish genotypes (Prevost & Wilkinson 1999). The values of the PIC and RP are comparable to those obtained with dominant markers in other species, e.g. *Podophyllum hexandrum* (Naik et al. 2010), *Jatropha curcas* (Grativol et al. 2011) and *Pongamia pinnata* (Sharma et al. 2014).

### Genetic diversity and differentiation

In order to estimate the genetic diversity of *B. serrata* Roxb. population, we applied both; band-based and allele frequency-based approach with non-Bayesian and Bayesian statistics so that the under-estimation or over-estimation of the diversity measures caused by the dominant marker systems could be avoided. A significant correlation between diversity measures i.e. P%, GD, I and Hs; with slight differences among them for different regions, confirms almost equal heterogeneity of the species population. The gene diversity of the sampled population has been found slightly higher than that reported by Nybom (2004) for out-crossing species (0.18). Nevertheless, Holsinger & Wallace (2007) suggested avoiding the comparison among the diversity measures resulted by the neutral markers (AFLP/RAPD) as these markers only show the differences in rates of mutations at their loci rather than reflecting different migration rates. However, there are few studies based on the tree species of the central Indian regions *viz.* Wang et al. (2011) investigated the genetic diversity of *D. sissoo* sampled from Indian regions and found 89.11% of polymorphism, 0.27±0.16 GD and 0.41±0.23 I. Ansari et al. (2012) sampled *Tectona grandis* L. f. from all along the central and peninsular Indian regions and found 80.30% polymorphism, 0.32 GD and 0.45 I. The comparison of the results from present investigation with above studies indicates that the species natural distribution and the level of genetic diversity exhibited are somehow equally influenced by the geographical conditions and the landscapes. The significant correlation found between genetic and geographical distances of the genotypes in our result also supports the fact.

The genetic structure of a population reflects the long-term evolutionary history of the species along with its mating system (Duminil et al. 2007), reproductive biology and gene flow influenced by the adaptive selection and fragmentation (Slatkin 1987, Nybom & Bartish 2000). In our investigation, the results of Bayesian model-based investigation through HICKORY found full-model as the most suitable for our data determining the influence of inbreeding within the populations. The low level of panmictic heterozygosity (Hs) for different regions and for overall population also supports it. Like other out-crossing species (Hamrick & Godt 1989; Nybom 2004) the *B. serrata* Roxb also exhibited higher within-population variability than among populations but the genetic variation within the population resulted by AMOVA was found comparatively lower than those reported in other out-crossing forest species (Nybom 2004, Wang et al. 2011, Ansari et al. 2012). A limited gene flow in *B. serrata* Roxb. is one of the important factors that have led to the high differentiation among sampled populations and the genotypes. The species has an entomophilous mode of pollination and the self-incompatible flowers support only cross-pollinated pollen to grow the pollen tubes. Moreover, only up to 10% fruit set has been observed in open-pollination condition (Sunnichan et al. 2005). In our field observation also, we noticed earlier abscission of unripe fruits and ripen fruit with empty seeds led to the total absence of natural regeneration in all sites of sampling. These bottlenecks contribute to genetic drift and signatory inbreeding depression to the species population reflected through different estimates.

The genetic differentiation measures G_ST_ and G_ST_-B have resulted in comparatively equal values indicating the cautions taken to employ the dominant markers for the investigation could be equivalent to the Bayesian model correction. The θ-statistics measures have exhibited differences in their values due to a low number of populations sampled for the investigation (Holsinger & Wallace 2004). The θ-I is supposed to be equivalent to Wright’s F_ST_ (Wright 1969) and its higher value (0.50±0.04) is expected because it results in the scaled allele frequency measured across the evolutionary time. The θ-II is a more appropriate measure for a small number of admixed population and its higher value (0.45±0.004) reflects the problematic gene exchange among the genotypes leading to high differentiation.

Among the Bayesian clusterings resulted by STRUCTURE, K=2 has been found the most suitable for our data on the basis of deltaK value. Nevertheless, the presence of inbreeding in sampled population and deviation from the HWE suggest preferring the dendrograph and the PCoA over the two admixed cryptic populations resulted by the program STRUCTURE because of its prior assumption of HWE. In PCoA, We have found Sarguja population closely related to Dhamtari population and the Narayanpur population has represented itself distinctly even after geographical closeness to Dhamatri. It might be due to the gene flow among these populations follows geographical limits along with their own biological constraints. On the other hand, the Narayanpur population might represent a different source of genetic origin of southern Indian region that could not be sampled. An almost equal polymorphism, genetic diversity, and heterozygosity but with highly differentiable genetic resource resulting to the distinctly different clusters indicate that the sampled populations may belong to a large metapopulation of the species that has been fragmented with due course of time due to its own intrinsic inability to cope up with natural and anthropogenic pressures later leading to geographical isolation.

*B. serrata* Roxb. is a highly out-crossing species with self-incompatibility and therefore, its own constraint to exchange the genes and the fragmentation of the population due to geographical isolation and commercial harvesting might be the reason for detected inbreeding pressure. The estimates of genetic diversity measures, genetic structure, and level of differentiation of a species population depend on the molecular marker system employed for the investigation. In our investigation, the dominant markers might be a reason behind low to moderate estimates of gene diversity, panmictic heterozygosity, and within-population variability. On the other hand, the Bayesian model-based analysis has given a higher value of θ-II because the θ-statistics has been developed specifically for the dominant markers (Spiegelhalter et al. 2002) and has been found more appropriate than G_ST_ and G_ST_-B to estimate the inbreeding and population differentiation (Holsinger et al. 2002, Holsinger & Wallace 2004). The comparatively higher variation in morphometric traits also indicates the existence of higher variability in the population. The northernmost region Sarguja and the southernmost region Narayanpur might represent different centers of genetic origin of a large metapopulation of the species and the genes from Sarguja are immigrating to the central region Dhamtari. For a better understanding of the influence of landscape and other geographical features on distribution and gene diversity of *B. serrata* Roxb. an investigation can be conducted with a larger number of locations representing its entire range of natural distribution applying co-dominant multiallelic marker system to cross verify the results of our investigation.

## CONCLUSION

*B. serrata* Roxb. has been over-exploited due to its significant quality of wood for pulp and paper industries. The technology in paper industries to use the species as a raw material has become obsolete and nowadays the oleo-gum-resin obtained from its wood is of significant medicinal value to heal arthritis and asthma. The species is endemic to India and unfortunately, there is not a single source of sustainable supply for the raw material required in pharmaceutical industries. Apart from all, the species is understudied and has not received much scientific attention to report on population genetic. Through the present investigation, we report that the available genetic resource of the species in central Indian region maintains high variation in wood fiber dimension and exhibits higher genetic variability within and among its sub-populations. In order to maintain the valuable forest genetic resource, the species need prior attention for the conservation. The locations of its natural distribution can be developed as strict nature reserves. On the other hand, a genetic improvement program should be initiated for selection and multiplication of elite genotypes. There is need to develop protocols for the artificial generation of the species through clonal and tissue culture techniques.

## ACKNOWLEDGEMENT

Authors are grateful to Director, Tropical Forest Research Institute, Jabalpur for Institutional support and to Director General, Indian Council of Forestry Research and Education, Dehradun for the financial support received under the project ID-175/TFRI/2011/Gen-4 (23).

